# Primary human lung fibroblasts exhibit trigger- but not disease-specific cellular senescence and impair alveolar epithelial cell progenitor function

**DOI:** 10.1101/2023.07.24.550385

**Authors:** Nora Bramey, Maria Camila Melo-Narvaez, Fenja See, Beatriz Ballester-Lllobell, Carina Steinchen, Eshita Jain, Kathrin Hafner, Ali Önder Yildirim, Melanie Königshoff, Mareike Lehmann

**Affiliations:** Comprehensive Pneumology Center (CPC) with the CPC-M bioArchive and Institute of Lung Health and Immunity, Helmholtz Zentrum München, Member of the German Center for Lung Research (DZL), Munich, Germany; 2Departamento de Ciencias Biomédicas, Facultad de Ciencias de la Salud, Universidad Cardenal Herrera – CEU, CEU universities, Valencia, Spain; Division of Pulmonary, Allergy & Critical Care Medicine, University of Pittsburgh Medical Center, Pittsburgh, PA, United States; Institute for Lung Research, Philipps-University Marburg, Member of the German Center for Lung Research (DZL), Marburg, Germany

**Author notes:** Correspondence: Prof Dr Mareike Lehmann, Institute for Lung Research, Philipps University Marburg, Hans-Meerwein Str 2, 35043 Marburg.

## Abstract

Aging is the main risk factor for chronic lung diseases including idiopathic pulmonary fibrosis (IPF) and chronic obstructive pulmonary disease (COPD). Accordingly, hallmarks of aging such as cellular senescence are increased in different cell types such as fibroblasts in the lungs of these patients. However, whether the senescent phenotype of fibroblasts derived from IPF or COPD differs is still unknown. Therefore, we characterized senescence at baseline and after exposure to disease-relevant insults (H_2_O_2_, bleomycin, and TGF-β1) in cultured primary human lung fibroblasts (phLF) from control donors, IPF, or COPD patients. We found that phLF from different disease-origins have a low baseline senescence. H_2_O_2_ and bleomycin treatment induced a senescent phenotype in phLF whereas TGF-β1 only had a pro-fibrotic effect. Interestingly, we did not observe any differences in susceptibility to senescence induction in phLF based on disease origin. However, exposure to different stimuli resulted in different senescent programs in phLF. Moreover, senescent phLF reduced colony formation efficiency of alveolar epithelial progenitor cells. In conclusion, the senescent phenotype of phLF is mainly determined by the senescence inducer and impairs alveolar epithelial progenitor capacity *in vitro*.

## Introduction

Chronic respiratory diseases are the third leading cause of death globally (1). Those include chronic obstructive pulmonary disease (COPD) and interstitial lung diseases such as idiopathic pulmonary fibrosis (IPF) (1). COPD is an inflammatory disease (2) characterized by small airway remodeling, emphysema, and chronic bronchitis (3). The main risk factors are cigarette smoking and age but exposure to other air pollution or pathogens also contribute to COPD (3). In contrast to that, IPF is a progressive fibrosing disease of unknown cause (4,5). Furthermore, higher age and exposure to cigarette smoke are the main risk factor for IPF (4,5). Familial cases of IPF have mostly been linked to mutations in genes encoding surfactant protein C and A2 (SFTPC, SFTPA2) and telomerases (TERT and TERC), that ultimately lead to telomere shortening, cellular senescence, and exhaustion of lung stem cells (4–7). Notably, incidence rates for both COPD and IPF increase in the elderly population (8). Different cellular hallmarks of aging have been described, including cellular senescence, which is characterized by an irreversible arrest of the cell cycle, a resistance to apoptosis, and a change in gene expression, including distinctive changes in protein secretion known as the senescence associated secretory phenotype (SASP) (9). All these features of cellular senescence are increased in the lung tissue from patients with IPF and COPD (10–13).

Fibroblasts are effector cells in both diseases, causing impaired tissue structure by aberrant deposition of extracellular matrix (ECM) on the one hand, and demonstrating hallmarks of senescence on the other hand (11,12,14–16). Recent progress in single cell omics revealed the existence of disease-specific cellular subtypes in the mesenchymal compartment (17–21). Although accumulation of senescent cells has been shown in both COPD and IPF, it remains unclear how senescence is induced in specific cell types and whether certain subtypes are more prone to different senescence stimuli. Therefore, here we used well-known senescence inducers to study susceptibility and senescence programs in primary human lung fibroblasts (phLF) from control donors, IPF, and COPD patients. In this study, we show that phLF from different disease origins have a low baseline senescence in culture. Moreover, stimuli and not disease origin determines the senescence phenotype in phLF. Finally, senescent fibroblasts modulate progenitor capacity of alveolar epithelial progenitors *in vitro*.

## Results

### Primary human lung fibroblast isolated from different lung diseases exhibit similar senescence phenotypes at baseline

The phLF derived from Donor, COPD or IPF patients were characterized at baseline for multiple well-accepted senescence markers after short (Day3) and prolonged (Day7) culture (Fig. 1A). First, we evaluated the senescence-associated-β (SA-β)-galactosidase activity, and observed a low percentage of SA-β-galactosidase+ cells in all three groups with no significant differences among them (Fig. 1B). Similarly, we did not observe any significant difference in proliferation between the 3 groups (Fig. 1C). We then determined the gene expression of the cyclin dependent kinase inhibitor 1A (*CDKN1A/P21*) and 2A (*CDKN2A/P16*) and the tumor suppressor protein 53 (*TP53*). For all conditions, we observed a stable expression over time with no statistical difference among the different origins (Fig 1D). Finally, we found no significant differences in the secretion of the SASP components: Plasminogen Activator Inhibitor 1 (PAI-1), Matrix Metalloproteinase-3 (MMP-3), GDF-15, or Interleukin 6 (IL-6), by fibroblasts from different disease origins (Fig. 1E). In conclusion, we observed a low level of senescence-related markers in cultured human primary fibroblasts irrespective of their disease origin.

**Figure 1.**
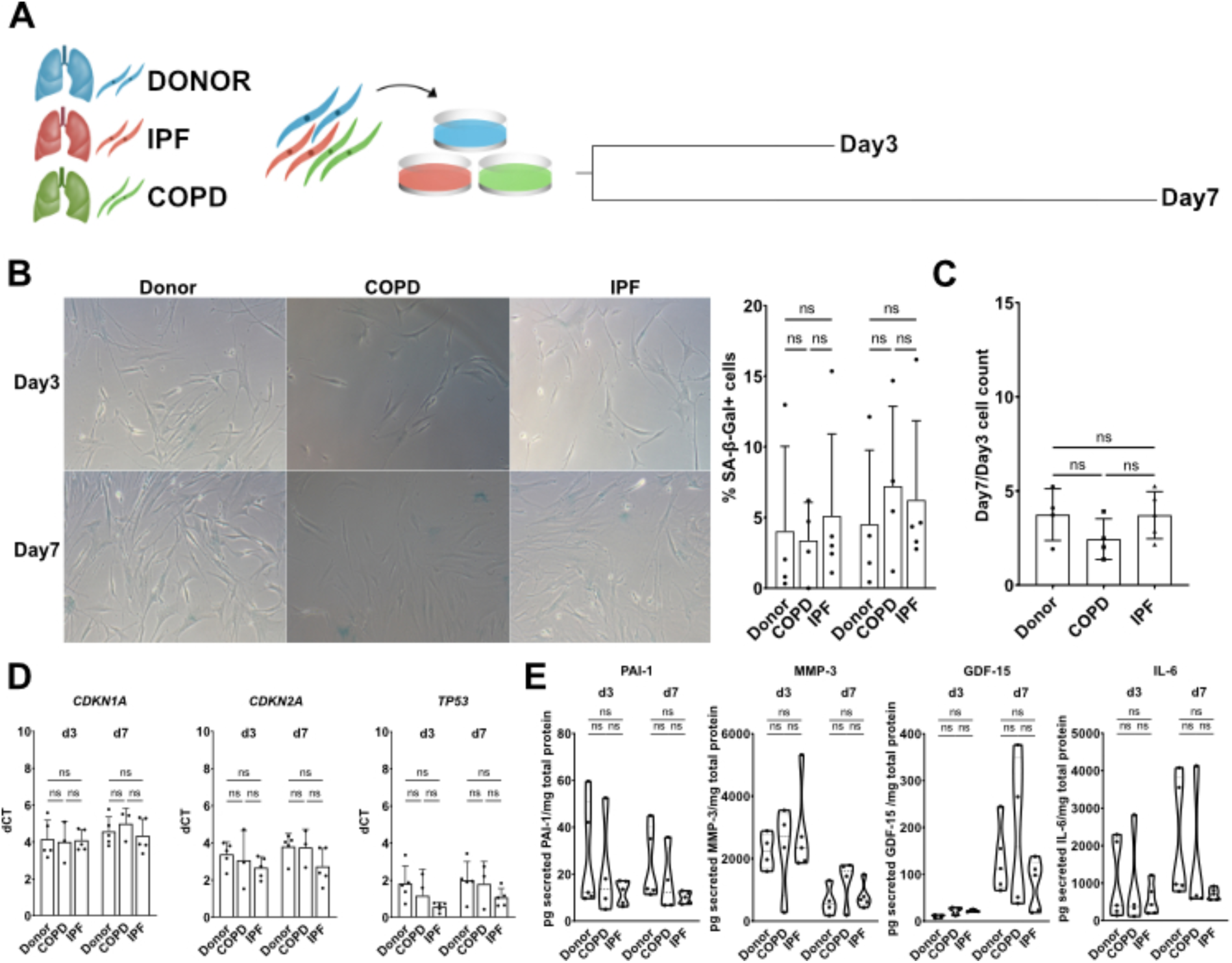
Primary human fibroblasts from different disease states do not show differences in senescence at baseline. A. Experimental design to characterize senescence in primary human fibroblasts from Donor, COPD, and IPF patients after short (Day3, d3) and prolonged (Day7, d7) culture. B. Representative images of SA-β-galactosidase staining of phLF from Donor, COPD, and IPF. C. Quantification of SA-β-galactosidase+ cells and proliferation rate calculated based on cell number are shown. Data points represent an average of 3 different regions of interest of at least 3 different biological replicates. D. qRT-PCR to assess gene expression of senescence-related markers (*CDKN1A, TP53, CDKN2A*) after 3 or 7 days in culture in phLF from different origins. E. ELISA of phLF supernatants that were cultured for 3 or 7 days. Data points represent different biological replicates of the concentration of each secreted protein (pg/ml) normalized to total cell protein content (mg/ml). All p-values (<0.05) were calculated based on Kruskal-Wallis test.

### Induction of senescence in primary human fibroblasts with different stimuli

Aging and exposure to cigarette smoke are the main risk factors for IPF and COPD and have been linked to increased stress-induced senescence. Therefore, we exposed pHLF to hydrogen peroxide (H_2_O_2_) or bleomycin, since they induce the release of reactive oxygen species (ROS) and genomic DNA-damage. Moreover, since previous studies showed that transforming growth factor beta 1 (TGF-β1), a well-known profibrotic mediator, not only promotes fibroblast activation but also senescence, we included this as a third senescence trigger (22). For bleomycin and H_2_O_2_, we observed a dose-dependent induction of SA-β-galactosidase activity (Fig. 2A, B) and used a dose of 3.3mU/ml for bleomycin and 180uM for H_2_O_2_, respectively, for further experiments, since these doses induced a high percentage of senescent cells (58.8% and 61.5%, respectively) and a significant reduction in cell number (Fig. 2C). Next, we tested whether single (S) or repetitive (R) treatment would induce different responses in senescence-related markers. We found that *CDKN1A/P21* was significantly induced (Fig. 2D) by repetitive treatment and therefore, continued with this treatment scheme for further experiments.

**Figure 2.**
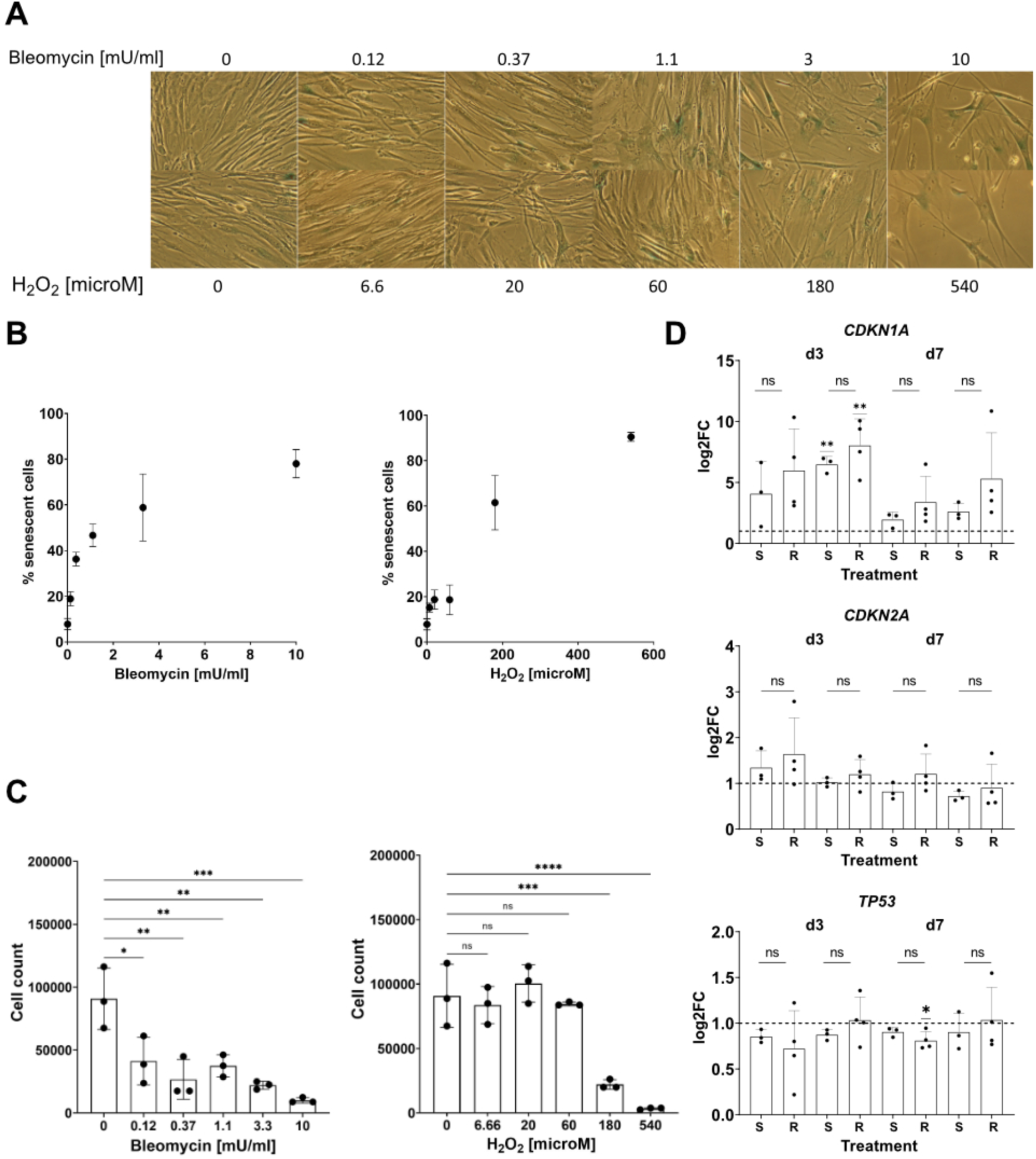
Establishment of treatment regimen for primary human fibroblasts to induce senescence. A. Representative images of SA-β-galactosidase staining after treatment of phLF with different concentrations of bleomycin and H_2_O_2_ to determine the best effective dose after 3 days of treatment. B. Titration of H_2_O_2_ and bleomycin concentration based on percentage of SA-β-galactosidase+ cells. C. Cell counts from primary human fibroblasts treated with different doses of bleomycin and H_2_O_2_. Single points represent replicates. *p-value<0.05 after one-way ANOVA test. D. qRT-PCR to assess gene expression of senescence-related markers after single or repetitive hit treatment regimens for 3 and 7 days. Single vs. repetitive hit: *p-value<0.05: Kruskal-Wallis test followed by Dunńs multiple comparisons test. Log2FC to Ctrl: *p-value<0.05: One sample t test.

Next, we addressed the senescence phenotype of phLF upon senescence induction with the chosen regimen and doses for 3 and 7 days (Fig. 3A). We analyzed several lines of donor or disease derived phLF (Table 1). PhLF treated with H_2_O_2_ and bleomycin showed significantly increased SA-β-galactosidase activity and decreased replication, consistent with a senescent phenotype (Fig. 3B, C). Notably, TGF-β1 treatment increased classical fibrotic markers (Suppl. Fig. 1) but did not affect SA-β-galactosidase activity or replication (Fig. 3B, C). The expression of *CDKN1A/P21* was increased by all treatments, while *CDKN2A/P16* was only induced by H_2_O_2_ and TGF-β1 treatment (Fig. 3D). Finally, we characterized the secretion of different SASP factors. Here, we found that H_2_O_2_ and bleomycin induced secretion of GDF-15 and MMP-3 and reduced PAI-1 secretion (Fig. 3E). Conversely, TGF-β1 treatment significantly induced IL-6 and PAI-1 secretion and reduced MMP-3 secretion (Fig. 3E). In conclusion, H_2_O_2_ and bleomycin induced a senescent phenotype in phLF characterized by cell cycle inhibition, reduced proliferation, SA-β-galactosidase activity, and secretion of SASP-related proteins. On the other hand, TGF-β1 treatment only showed an effect on *CDKN1A/P21* and *CDKN2A/P16* expression together with secretion of well-known downstream mediators of the TGF-β1 signaling pathway like PAI-1 and IL-6.

**Figure 3.**
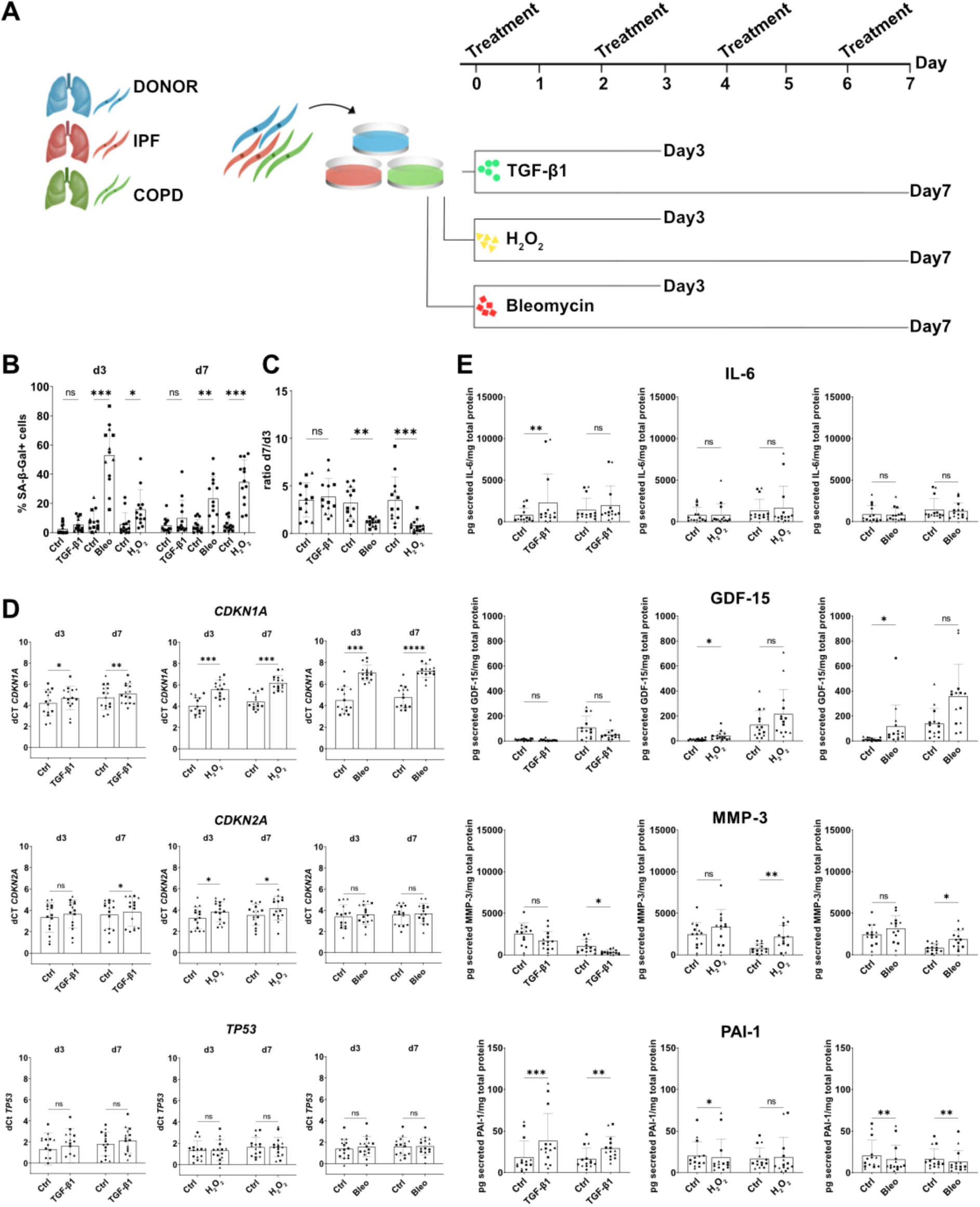
Induction of senescence in primary human fibroblasts with different stimuli. A. Experimental design to characterize senescence induction on primary human fibroblasts from Donor, COPD, or IPF patients after treatment with TGF-β1, H_2_O_2_, or bleomycin, respectively. B. Quantification of SA-β-galactosidase activity after 3 and 7 days after treatment. C. Proliferation rate (Cell count d7/d3) after treatment with H_2_O_2_, bleomycin, and TGF-β 1. Data points represent biological replicates from donor (square), IPF (circle), and COPD (triangle). *p-value<0.05: Kruskal-Wallis test followed by Dunńs multiple comparisons test. D. qRT-PCR to assess gene expression of senescence-related markers after treatment with H_2_O_2_, bleomycin, and TGF-β1 E. ELISA of supernatants of primary human fibroblasts treated with H_2_O_2_, bleomycin, and TGF-β 1. Data points represent different biological replicates from donor (square), IPF (circle), and COPD (triangle) of the concentration of each secreted protein (pg/ml) normalized to total lysate protein content (mg/ml). *p-value<0.05 based on paired-Friedman T test.

**Table 1.** Patient demographics and clinical data.

| Gender | Age (Mean+SD) | Age | Smoker | Diagnosis |
| --- | --- | --- | --- | --- |
| female | 70+/-11.2 | 72 | ex-smoker >6 months | Lung cancer/Peritumor |
| female |  | 67 | smoker | Lung cancer/Peritumor |
| male |  | 84 | ex-smoker >6 months | Lung cancer/Peritumor |
| female |  | 57 | NA |  |
| female | 63.6+/-8.8 | 56 | ex-smoker | IPF, no PAH |
| male |  | 72 | unknown | IPF, no PAH |
| female |  | 54 | never smoker | ILD collagenosis, no PAH |
| male |  | 73 | unknown | IPF, no PAH |
| male |  | 63 | ex-smoker >6 months | IPF, precapillary PH |
| male | 67.7+/-4.96 | 67 | ex-smoker >6 months | COPDIV |
| male |  | 73 | ex-smoker >6 months | COPDIV; no PAH |
| female |  | 69 | ex-smoker | COPDIV; low PAH |
| male |  | 62 | ex-smoker | COPDIV; Emphysema |
| female |  |  | NA | COPDIV |

### PhLF isolated from different lung disease exhibit no differences in inducibility of senescence

Next, we addressed whether the susceptibility to cellular senescence differs depending on the different disease backgrounds of phLF. We compared the level of senescence induction in donor-, IPF-, or COPD-derived phLF. First, we calculated the fold change to control of SA-β-galactosidase+ cells for each group at day 3 and day 7, to evaluate whether the effect differed among disease origins. Here, we found no significant differences among disease origins (Fig. 4A, B). Then, we determined the expression of three cell cycle regulators. Interestingly, we did not observe any significant difference in gene expression of *CDKN1A/P21* and *CDKN2A/P16* or *TP53* among the different disease origins (Fig. 4C). In conclusion, phLF from different disease origins have similar susceptibility to the tested senescence inducers.

**Figure 4.**
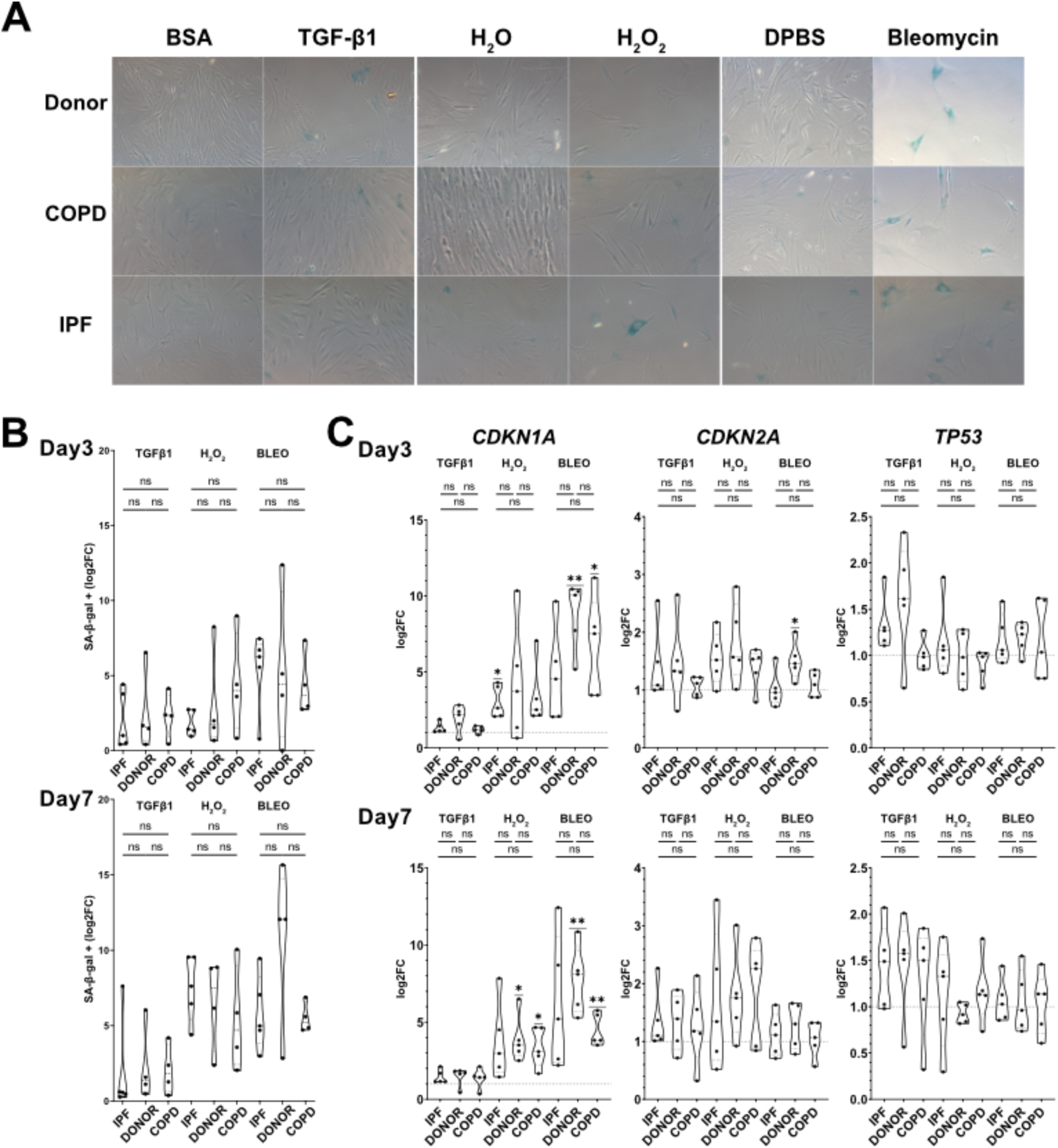
No differences in senescence inducibility based on cellular origin. A. Representative images of senescence inducibility assay based on SA-β-galactosidase activity after 7 days of treatment with bleomycin, H_2_O_2_, or TGF-β1 in primary human fibroblasts from Donor, COPD, and IPF patients. B. Quantification of SA-β-galactosidase activity in primary human fibroblasts from Donor, COPD, and IPF patients after 3 and 7 days of treatment with H_2_O_2_, bleomycin, or TGF-β1. C. qRT-PCR to assess gene expression of senescence-related genes (CDKN1A, CDKN2A, TP53) by in primary human fibroblasts from Donor, COPD, and IPF patients after 3 and 7 days of treatment with H_2_O_2_, bleomycin, or TGF-β1. Origin: *p-value<0.05: Kruskal-Wallis test followed by Dunńs multiple comparisons test. Log2FC to Ctrl: *p-value<0.05: One sample t test.

### Different stimuli induce different senescence programs

We next compared the senescence signatures of phLF induced by TGF-β1, bleomycin, and H_2_O_2_. For all origins and treatments, we observed that SA-β-galactosidase activity increased over time (Fig. 5A). Interestingly, we observed that H_2_O_2_ induced a significant increase only in fibroblasts derived from IPF patients, whereas bleomycin significantly induced SA-β-galactosidase activity in both Donor- and COPD-derived fibroblasts after 7 days (Fig. 5A). Again, TGF-β1 did not induce SA-β-galactosidase activity in any treatment or cell origin (Fig. 5A). As observed before, H_2_O_2_ and bleomycin strongly induced *CDKN1A/P21* expression, whereas *CDKN2A/P16* and *TP53* had a more moderate increase over time (Fig. 5B). Moreover, H_2_O_2_ and bleomycin induced the expression of *PAI-1*, *ACTA2*, and Fibronectin 1 (*FN-1*) after 7 days of culture (Fig. 5B). Conversely, TGF-β1 did not increase senescence-related genes but had a stronger effect on the expression of pro-fibrotic markers such as *ACTA2*, *FN-1*, and Collagen 1 (*COL1A1*) (Fig. 5B). Finally, SASP profiles were more similar between H_2_O_2_ and bleomycin as shown by an initial induction of IL-6, and GDF-15 followed by later induction of MMP-3 secretion (Fig. 5C). TGF-β1 only induced the secretion of PAI-1 and IL-6 (Fig. 5C). In conclusion, we observed specific senescence programs depending mainly on the trigger.

**Figure 5.**
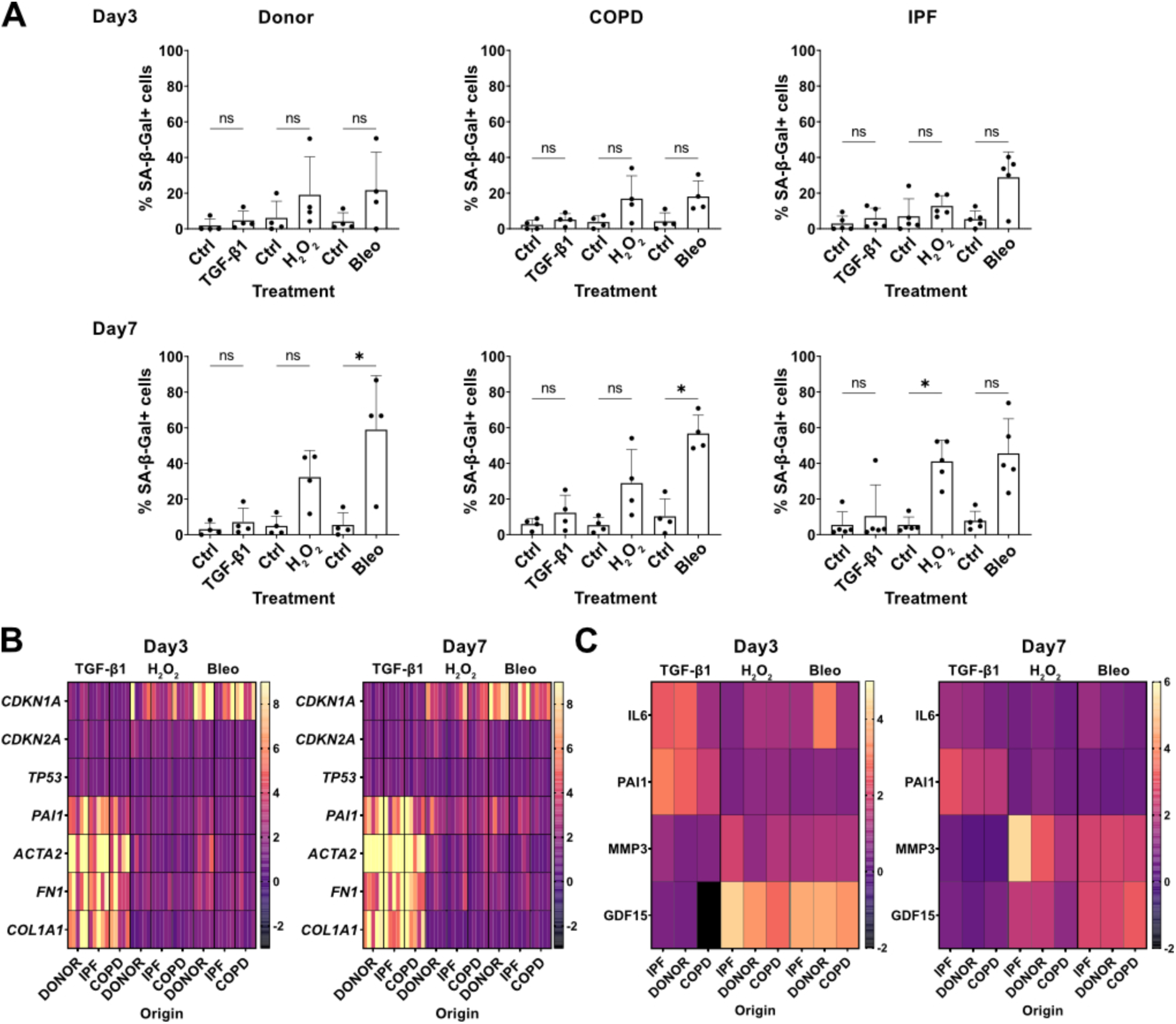
Different stimuli induce different senescence program. A. Quantification of SA-β-galactosidase activity induction in primary human fibroblasts from Donor, IPF and COPD after treatment with H_2_O_2_, bleomycin and TGF-β1 for 3 and 7 days. *p-value<0.05: Kruskal-Wallis test followed by Dunńs multiple comparisons test. B. Heatmap of qRT-PCR to assess gene expression of senescence-(*CDKN1A, CDKN2A, and TP53*) and fibrosis-(*ACTA2, PA-1, FN-1, COL1A1*) related genes after treatment with H_2_O_2_, bleomycin or TGF-β1 for 3 and 7 days. Single rows represent biological replicates from Donor-, IPF-, and COPD-derived primary human fibroblasts. C. Heatmap of the SASP of Donor-, IPF-, and COPD-derived primary human fibroblasts as assessed by ELISA after treatment with H_2_O_2_, bleomycin or TGF-β1 for 3 and 7 days. Single rows represent the average expression of at least 4 biological replicates. Concentration of each secreted protein (pg/ml) normalized to total lysate protein content (mg/ml).

### Senescent fibroblasts disrupt progenitor potential of lung alveolarvepithelial cells

Senescent fibroblasts accumulate in COPD and IPF lungs. In the alveolar niche the epithelial and mesenchymal compartments closely interact but the paracrine effects of senescent cells in this environment are still understudied. To explore this, we tested whether senescent fibroblasts influence stem cell potential of distal lung progenitor cells by co-culturing them in an organoid assay (Fig. 6A). After 14 days, we observed formation of both alveolar (small and dark) and bronchiolar organoids (big with a lumen) in all conditions (Fig. 6C). However, phLF from donor patients treated with bleomycin and H_2_O_2_ significantly reduced colony formation efficiency (CFE) (Fig. 6B). Moreover, epithelial progenitors co-cultured with bleomycin-treated phLF formed significantly smaller organoids than controls (Fig. 6B). Finally, to characterize the cellular composition of the formed organoids, we stained them for surfactant protein C (SP-C), a marker for alveolar type 2 (AT2) cells, Keratin-8 (Krt8), a transdifferentiation marker for AT2 cells, and Acetylated Tubulin (ACT), a marker for airway epithelium as shown in figure 6C. In conclusion, the co-culture with senescent fibroblasts reduced stem cell capacity of alveolar epithelial progenitor cells *in vitro*.

**Figure 6.**
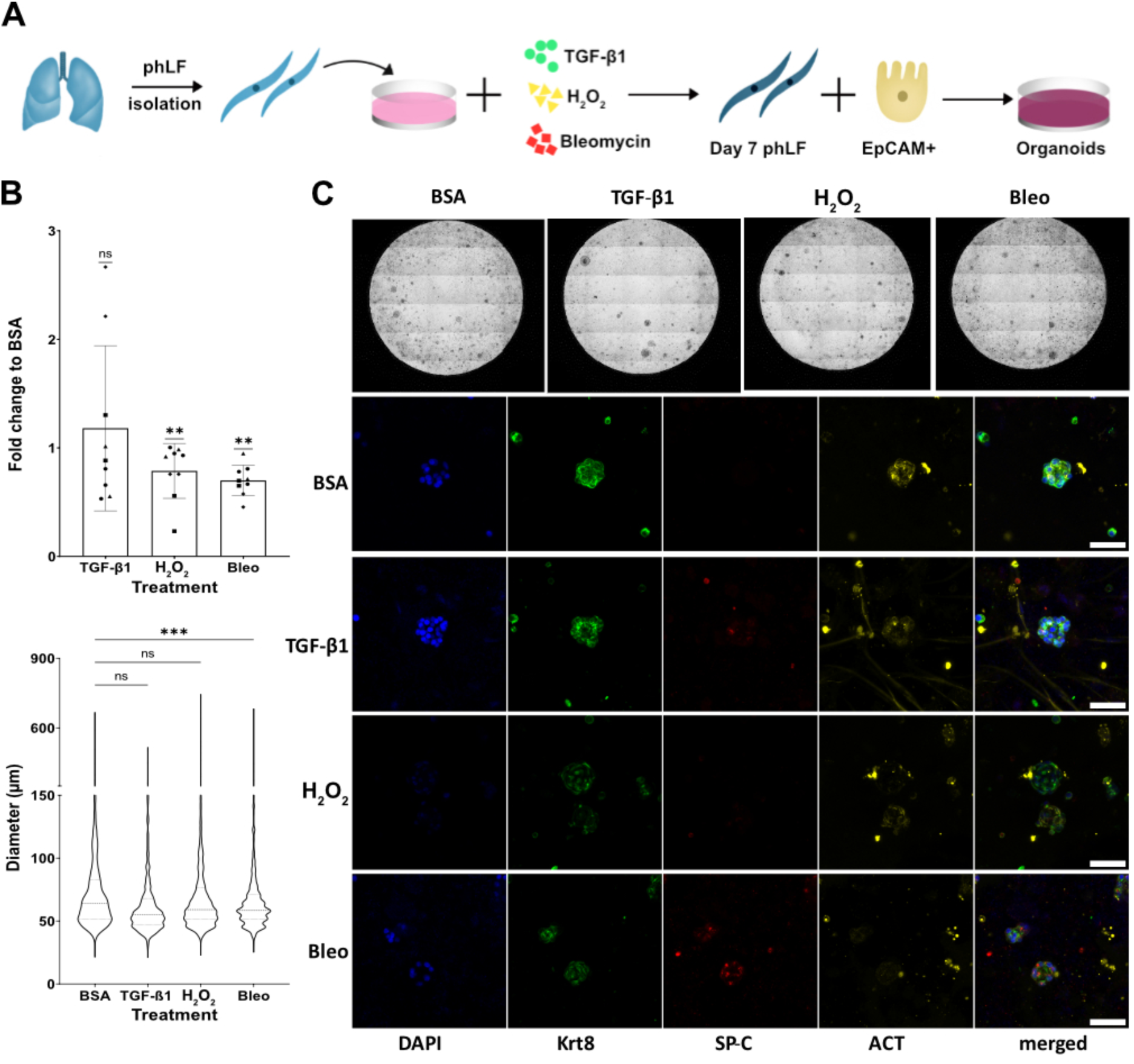
Senescent fibroblasts disrupt progenitor potential of lung epithelial cells. A. Experimental design. Primary mouse lung epithelial cells were co-cultured with primary human fibroblasts (Pre-treated with H_2_O_2_, bleomycin or TGF-β1 for 7 days) for 14 days. B. Fold change to BSA control for colony formation efficiency (CFE, top) and average spheroid size quantification (bottom). Single points represent 4 biological replicates (marked by shape) with 2 technical replicates for each. *p-value<0.05 based on One-sample t-test (CFE) or Kruskal-Wallis test (size). B. Representative bright-field images of whole wells and single organoids stained to evaluate the expression of surfactant protein C (SP-C), Acetylated Tubulin (ACT), and Keratin-8 (Krt8).

## Discussion

Aging is the main risk factor for CLDs such as COPD and IPF and previous studies have shown that senescent cells accumulate with age (23). Although the mechanism is not fully understood, senescent cells can evade immune clearance, thereby accumulating and promoting organ dysfunction (24–27). Indeed, elevated levels of *CDKN1A/P21*, *CDKN2A/P16,* and SA-β-galactosidase were found in fibroblasts from both IPF and COPD lungs and therefore, have been linked to the disease pathobiology (16,23,28–30). However, whether the senescent phenotype is different depending on the disease background is not well understood. Therefore, in this study we aimed to characterize the senescence of phLF from control, IPF, and COPD patients at baseline and after exposure to different senescence inducers. Finally, we used an organoid assay to study the crosstalk between epithelial and mesenchymal cells *in vitro*.

First, we characterized classical markers of senescence in phLF at baseline. Here, we found that fibroblast from control, IPF, and COPD had a very similar phenotype characterized by low expression of senescence-related markers: *CDKN1A/P21*, *CDKN2A/P16* and *TP53* gene expression, SA-β-galactosidase activity, and low secretion of SASP-related components. However, all these markers increased with prolonged culture as described for replication-induced senescence (31). Previous studies showed that fibroblasts originated from COPD patients had an elevated senescence signature as judged by enhanced expression of *P21*/*CDKN1A* and *CDKN2A/P16*, increased SA-β-galactosidase activity (32,33), reduced proliferation rates (30,33) and higher secreted levels of proteins associated with the SASP (15). Moreover, they inhibited canonical WNT-β-catenin signaling in alveolar epithelial cells by secreting WNT-5A, leading to stem cell exhaustion and impaired lung repair (34). Similarly, phLF obtained from IPF lung tissue, showed decreased proliferation rates, increased expression of *CDKN1A/P21*, *CDKN2A/P16*, and *TP53* as well as senescence-related morphological changes (16). The fact that we did not observe any major differences in senescence markers among disease origin, as reported previously (16,30,32,33), could be explained by different isolation protocols. For example, in this study we isolated phLF by enzymatic digestion contrary to outgrowth from tissue pieces that other studies used (32). The composition of the isolated phLF can also vary depending on different anatomical localizations such as airway (33) versus whole lung (16,30). Moreover, characteristics of different donor fibroblasts including smoking status, passage number and different culturing conditions or supplements of the culture media such as FCS could influence senescence readouts (35). Finally, all these isolation methods do not distinguish among different fibroblast subtypes described for the lung. In the past decades, it was believed that ACTA2+ positive myofibroblasts were the main contributor for ECM deposition in the IPF lung (19). However, recent single cell based studies have revealed that IPF lungs have a higher heterogeneity in fibroblasts than control lungs and that these subpopulations coexist in the lung and might play different roles in the disease progression (17,19). Interestingly, the susceptibility to typical fibrotic and senescence inducers such as bleomycin, has been shown to differ among these fibroblast subpopulations in the mouse lung (19). Consequently, the response to the stimuli used in this and other studies might be defined by the composition of the isolated and treated fibroblast population. Therefore, the development of isolation protocols for primary lung fibroblasts based on the newly described markers would help to find specific disease-relevant cellular responses of these subpopulations that could be therapeutically targeted. Cellular senescence can be induced by several stimuli such as increased oxidative stress, caused by exposure to cigarette smoke, or genomic DNA damage, induced by chemotherapeutic agents. Therefore, we used H_2_O_2_ and bleomycin to mimic these insults in vitro. Moreover, we also included TGF-β1, since it has been shown to induce a senescent phenotype in phLF (16,22). In our study, only H_2_O_2_ and bleomycin induced cell cycle arrest as measured by *CDKN1A*/*CDKN2A* gene expression and other senescence-related markers such as reduced proliferation, increased SA-β-galactosidase activity, and increased secretion of GDF-15 and MMP-3 after 7 days. Notably, TGF-β1 only induced the expression of pro-fibrotic markers (*ACTA2*, *FN-1*, *COL1A1*, and *PAI-1*) as well as secretion of IL-6 and PAI-1 but did not induce a clear senescence phenotype. This could be explained by the different doses used in this study (5ng/ml) and the TGF-β1-induced senescence, in which higher doses were used (22). Previous studies showed that aged individuals have around 3-4 ng/ml circulating TGF-β1 in plasma (36). Therefore, based on results using a physiologically relevant dose, we propose TGF-β1 as a pro-fibrotic rather than a senescence stimulus.

Next, we addressed whether the susceptibility to senescence was different among the different disease origins. Here, we found that IPF fibroblasts had a trend towards a reduced response to all stimuli in comparison to Donor- and COPD-derived fibroblasts as previously described (37). However, as observed at baseline, we did not find any significant difference in the senescence response among the different disease origins. This could be explained by the treatment regimens, which consistently induce a senescent phenotype overriding the cell origin. In conclusion, we found that the gene expression and SASP profiles were more similar among bleomycin and H_2_O_2_ treatment than after TGF-β1, suggesting that the senescence response is mainly mediated by the specific trigger, in this case DNA damage and oxidative stress, than by cellular predispositions. Despite the similar pattern in the induction of senescence- and fibrosis-related markers for bleomycin and H_2_O_2_ treatment, we also observed differences in the effect size for the tested markers. For example, the gene expression of *CDKN2A* or the secretion of MMP-3 was more pronounced on H_2_O_2_-treated phLF. Therefore, more comprehensive analysis of gene expression changes and secreted factors might be useful to better understand differences among these two senescence inducers.

Senescent cells can modulate their microenvironment in a paracrine manner by their SASP or direct interaction (23,38). We assessed the secretion of proteins related to inflammation and ECM deposition after induction of senescence. Here, we found that bleomycin and H_2_O_2_ induced the secretion of GDF-15, MMP-3, and PAI-1. On the other hand, TGF-β1 treatment induced the secretion only of proteins downstream of it signaling pathway: IL-6 and PAI-1. In IPF lungs, MMP-3 is secreted by different cell types including fibroblasts and has been linked to lung epithelium dysfunction and poor regenerative capacity as well as fibroblasts activation (39,40). GDF-15 and PAI-1 are well known SASP factors that also have been linked to inflammation and ECM remodeling in the diseased lung (41–43). Interestingly, as previously described for paraquat-induced cellular senescence of lung fibroblasts (32), here we also observed that bleomycin and H_2_O_2_ decreased the expression of COL1A1 in phLF. Moreover, bleomycin and H_2_O_2_ induced the gene expression of *PAI-1* and *FN-1*. This suggests that senescent fibroblasts can contribute to ECM remodeling as seen in CLDs. Given the changes observed in ECM-related genes and SASP profiles of senescent phLF, we used an organoid assay to evaluate whether co-culture with them would alter stem cell function of alveolar progenitor cells. Here, we found that both bleomycin and H_2_O_2_-induced senescence significantly reduced progenitor cell capacity as assessed by colony forming efficiency. However, only bleomycin significantly altered the size of the formed organoids. These could be explained by differences in the SASP and ECM-related genes expression between bleomycin- and H_2_O_2_-induced senescence programs in phLF.

In conclusion, this study provides novel insights into the senescence phenotype of primary human lung fibroblasts exposed to disease-relevant insults. Further characterization of these phenotypes using state of art techniques such as single cell sequencing could help to elucidate the underlying mechanism that defines these senescent programs. Moreover, *in vitro* organoid assays allowed the characterization of cell-to-cell interactions that modulate the regenerative capacity of the lung progenitors. This could be complemented by other 3D models such as precision cut lung slices (PCLS) that would allow the study of the role of different senescent fibroblasts subpopulations in situ, since this model preserves the ECM and most cellular compartments that interact in the lung niche.

## Methods

### Primary human cells

Primary human lung fibroblasts derived from age-matched control, COPD, and IPF patients (Table 1) were obtained from the CPC-M bioArchive at the Comprehensive Pneumology Center (CPC Munich, Germany).

### Ethic statement

The study was approved by the local ethics committee of the Ludwig-Maximilians University of Munich, Germany (Ethic vote 19-630). Written informed consent was obtained for all study participants.

### Culture and harvesting of primary human lung fibroblasts

Primary human lung fibroblasts were isolated as previously described (44) derived from age-matched control, COPD, and IPF patients (53% male, mean age 66 years old) were cultivated in DMEM/F-12 (Life Technologies, USA) with 1% penicillin/streptomycin (10.000U/ml, Life Technologies, USA), and 10% Fetal Bovine Serum (PAN Biotech, Germany).

For qPCR, Western Blot and ELISA experiments, phLF were seeded on 6 well plates at a density of 4.5 x 10^4^ cells per well. After 24h of incubation, treatment solutions were applied and changed every 48h. After day3 and day7 of treatment, supernatants of cells designated for protein isolation were collected, centrifuged and frozen at -80°C. For RNA isolation, treatment solution was removed, the wells were washed twice with DPBS and cells were frozen directly at -80°C.

### Induction of cellular senescence

Primary human lung fibroblasts (passages 3-9) were exposed to 5ng/ml recombinant human TGF-β1 (R&D Systems, 240-B-002), 180μM H_2_O_2_ (Sigma, Germany) or 3,3mU/ml bleomycin sulfate (Sigma, Germany) in DMEM/F-12 with 1% Pen/Strep and 5% FBS. Negative control solutions contained equivalent volume of 0.1% BSA in PBS for TGF-β1, plain medium for H_2_O_2_ or DPBS for bleomycin.

### Organoid assay

Primary human fibroblasts were treated as described before. Then, phLF were treated with Mitomycin C (10 µg/ml) for 2h, at 37°C and 5% CO_2_ to stop proliferation. Then, phLF were washed with 1X DPBS and kept in fresh medium for at least 1h at 37°C and 5%CO_2_. Trypsin was used to obtain a single cell suspension that was kept on ice until use. Mouse lungs were flushed with DPBS, filled with Dispase and incubated in Dispase solution for 45 min. Lung tissue was manually minced and cells were filter using 100um and 40um nylon filters as previously described (45). Single cell suspension was then MACS sorted to isolated CD45-/CD31-/EpCAM+ cells. Human fibroblasts and murine AT2 cells were mixed 1:1 ratio (10.000 cells each), spun down for 10 min at 300 xg. The cell suspension was mixed in a 1:1 ratio with Matrigel loaded in 96-well plates. Plates were incubated at 37°C for 15 min and organoid medium supplemented with Rock inhibitor (Ri) was added on top. After 48h, Ri was removed and medium was changed every 2-3 days. After 14 days plates were imaged using a LifeCellImager Observer Z1 (Zeiss, Germany) at 5X. Maximum projections were generated on Zen software (Zeiss, Germany). Organoid size and number was determined using the Napari organoid counter as previously described (46,47).

### RNA isolation and RT-qPCR

RNA was isolated using the peqGOLD Total RNA Kit (VWR) according to manufacturer’s instructions. RNA concentration was quantified with a Nanodrop 1000 (Peqlab). For cDNA synthesis, RNA was diluted with RNAse free water to a concentration of 500ng/20ul and denatured at 70°C for 15min in a Mastercycler (Eppendorf). Mastermix was prepared by mixing: Random Hexamers (10uM, Invitrogen, cat.# 100026484), dNTP Mix (2mM, Thermo Scientific, cat.# R0192), 5x First-strand buffer (1X, Invitrogen, cat.# Y02321), 0.1 M DTT (40mM, Invitrogen, cat.# Y00147), RNAse Inhibitor (4U/ul, Applied Biosystems, cat.# N8080119), and M-MLV Reverse Transcriptase (10U/ul, Invitrogen, cat.# 28025013) was prepared and 20ul were added to each probe of denatured RNA before starting reverse transcription program (1 cycle at 20°C for 10min, 1 cycle at 43°C for 75min, 1 cycle at 99°C for 5min). The cDNA was then diluted 1:2-1:5 using RNAse free water and kept at -20°C until use. RT-qPCR mix was prepared using 1X Light Cycler 480 SYBR Green Master (Roche, Germany) and primer pairs at 5uM (Table 2). Then, samples were run in a Light Cycler 480II (Roche, Germany): 1 cycle at 50°C for 2 min, 1 cycle 95°C for 5 min, followed by 45 cycles of 1X 95°C for 5s, 1X 59°C for 5s, 1X72°C for 5s. The dCt values were determined by a two-derivative method and log2FC were calculated based on the 2^−ΔΔC^ method (48).

**Table 2.**
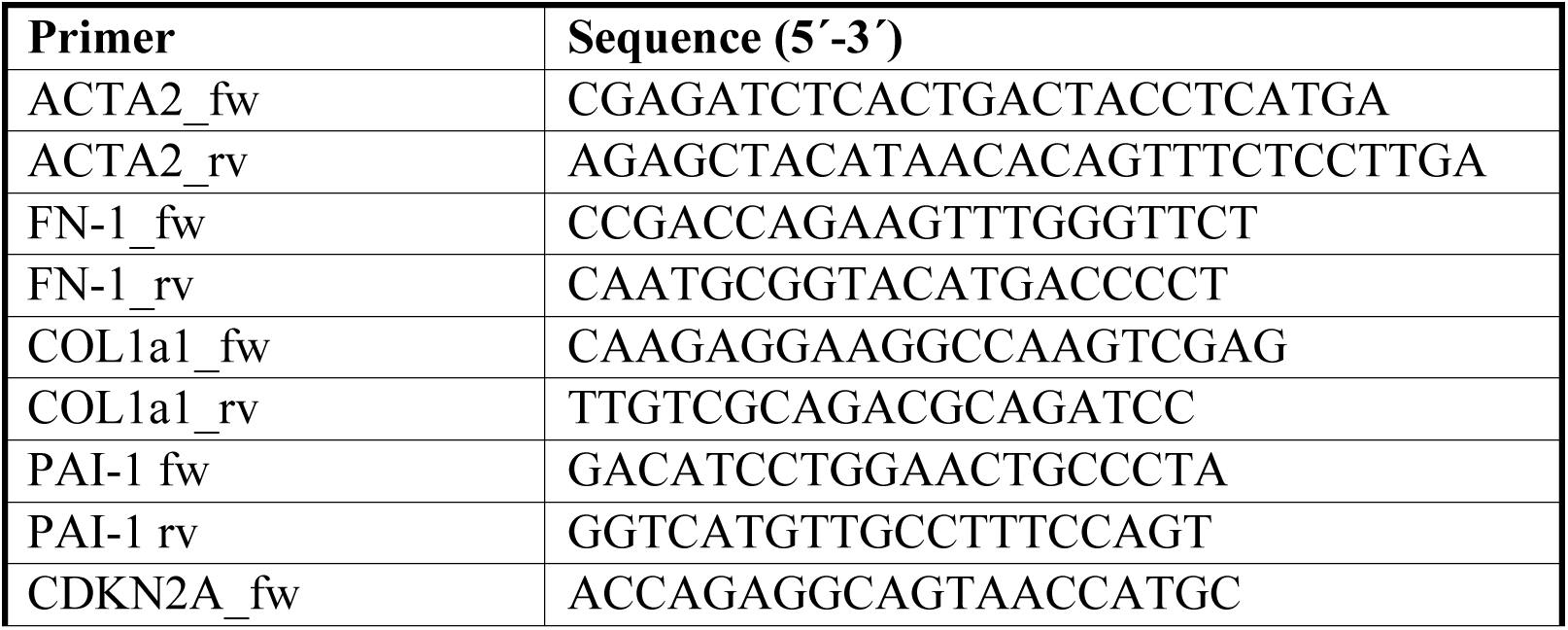
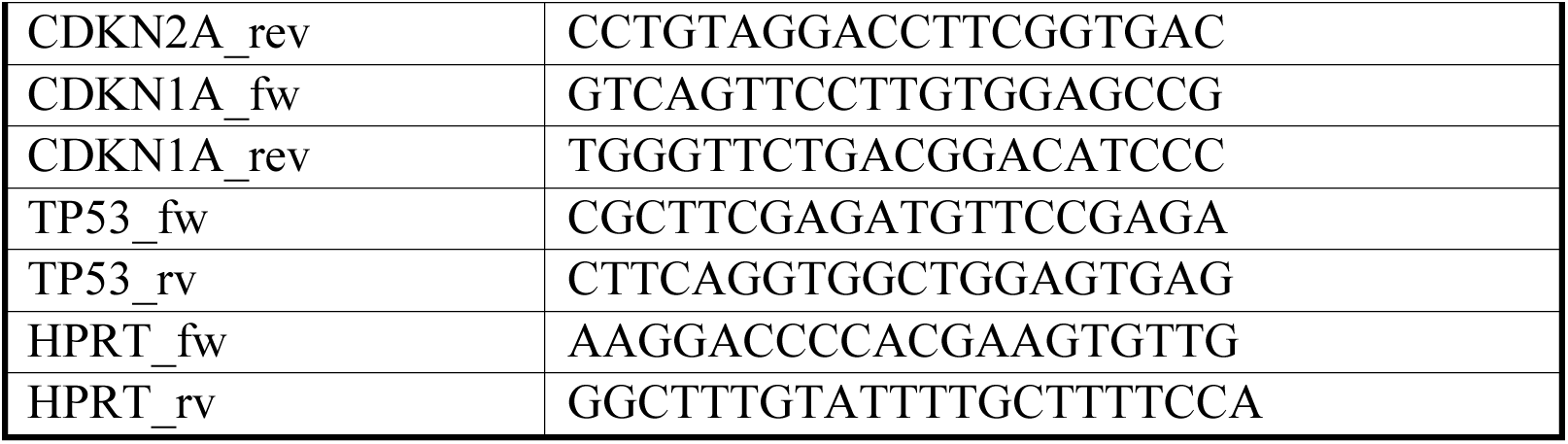
Primer list.

### ELISA

ELISA was performed using the human IL-6 Duo-Set (R&D Systems, cat.# DY206), human GDF-15 Duo Set (R&D Systems, cat.# DY957), human Serpin E1/PAI-1 Duo Set (R&D Systems, cat.# DY1786), and human total MMP-3 (R&D Systems, cat.# DY513) according to manufacturer’s instructions. Absorbance was measure at 450nm using the microplate reader Sunrise (Tecan GmbH, Germany). Final absorbance values were calculated by subtracting the background signal. Concentrations were calculated by interpolation of a linear regression based on the standard curve. Concentrations were normalized to total protein content of the cell lysate at the final collection time point. To generate the heatmap, a fold change to control (treatment/control) was calculated and the mean of at least 3 different biological replicates is shown.

### Senescence associated β-Galactosidase staining

For SA-β-galactosidase staining, phLF were seeded at a density of 8.0 x 10^3^ cells per well on a 12 well plate and senescence induction was performed as described above. Then, at collection points cells were washed twice with PBS. Then, cells were fixed and stained using the Senescence β-galactosidase staining Kit from Cell Signaling (cat. #9860) according to manufacturer’s instructions.

### Immunofluorescence staining

Organoids were fixed with ice-cold methanol for 10 min at -20°C, washed 3X with DPBS and kept in DPBS until staining. Then, organoids were blocked with 5\% donkey normal serum in 0.1% triton in 1X PBS (PBST) for 1h at RT. Then, primary antibodies (Table 3) were diluted in 1% normal donkey normal serum, added to the samples, and these were incubated at 4°C overnight. Samples were then washed 3X for 20 min with 0.1% PBST and secondary antibodies plus DAPI were added (Table 3). Samples were incubated for 2h at room temperature, plates were washed with 1X PBS and directly imaged using a LSM 710 Confocal microscope (Zeiss, Germany) at 10X for whole well images or 40X for single organoid images. Mean fluorescence intensity was quantified using ImageJ (Fiji) and data and plots were generated and analyzed in GraphPad Prism 9.5.1.

**Table 3.** Antibody list.

| Target protein | Host | Company | Ref. No |
| --- | --- | --- | --- |
| Actin, α-smooth muscle | mouse | sigma | A5228 |
| P21 [EPR18021] | rabbit | abcam | ab188224 |
| Phospho-histone H2A.X (Ser139) clone JBW301 | mouse | Millipore | 05-636 |
| P21 WAF1 (Ab-1)(EA10) | mouse | Merck | OP64 |
| p16 (JC8) | mouse | Santa Cruz | sc56330 |
| Ki67 | mouse | abcam | ab6526 |
| ACT | mouse | abcam | ab24610 |
| SP-C | rabbit | abcam | ab3786 |
| Krt8 | rat | DSHB | TROMAI |
| Anti-rat-488 | donkey | Invitrogen | A21208 |
| Anti-rabbit-647 | donkey | Invitrogen | A31573 |
| Anti-mouse-568 | donkey | Invitrogen | A10037 |
| Anti-mouse-555 | goat | Invitrogen | A21424 |
| Anti-rabbit-488 | goat | Invitrogen | A11008 |

### Data collection and analysis

Primary human fibroblasts from Donor, COPD, and IPF were used for different experiments in this study (Table 1). For titration (Fig. 2) and organoid assays (Fig. 6) phLF only from control donors were used and single points represent different biological or technical replicates as indicated in figure legends. To analyze the capacity of the triggers used to induce senescence (Fig. 3) we pooled together the data collected using all the different samples listed in Table 1. Here, single points represent biological replicates and points shape indicate the disease origin.

Finally, to study differences linked to the background disease we separated the samples listed in Table 1 into three different groups and used the data collected with these same samples for downstream analysis. Here, single points represent biological replicates and points shape indicate the disease origin.

## Acknowledgments

We gratefully acknowledge the provision of human biomaterial and clinical data from the CPC-M bioArchive and its partners at the Asklepios Biobank Gauting, the LMU Hospital and the Ludwig-Maximillians-Universität München. We thank the patients and their families for their support. ML acknowledges funding from the Deutsche Forschungsgemeinschaft (DFG, German Research Foundation) – 512453064, Federal Institute for Risk assessment (Bundesinstitut für Risikoforschung, BfR) 60-0102-01.P588, and the German Center for Lung Research (DZL).

