## Supplementary figure for "Primary human lung fibroblasts exhibit trigger- but not disease-specific cellular senescence and impair alveolar epithelial cell progenitor function"

**Supplementary figures**

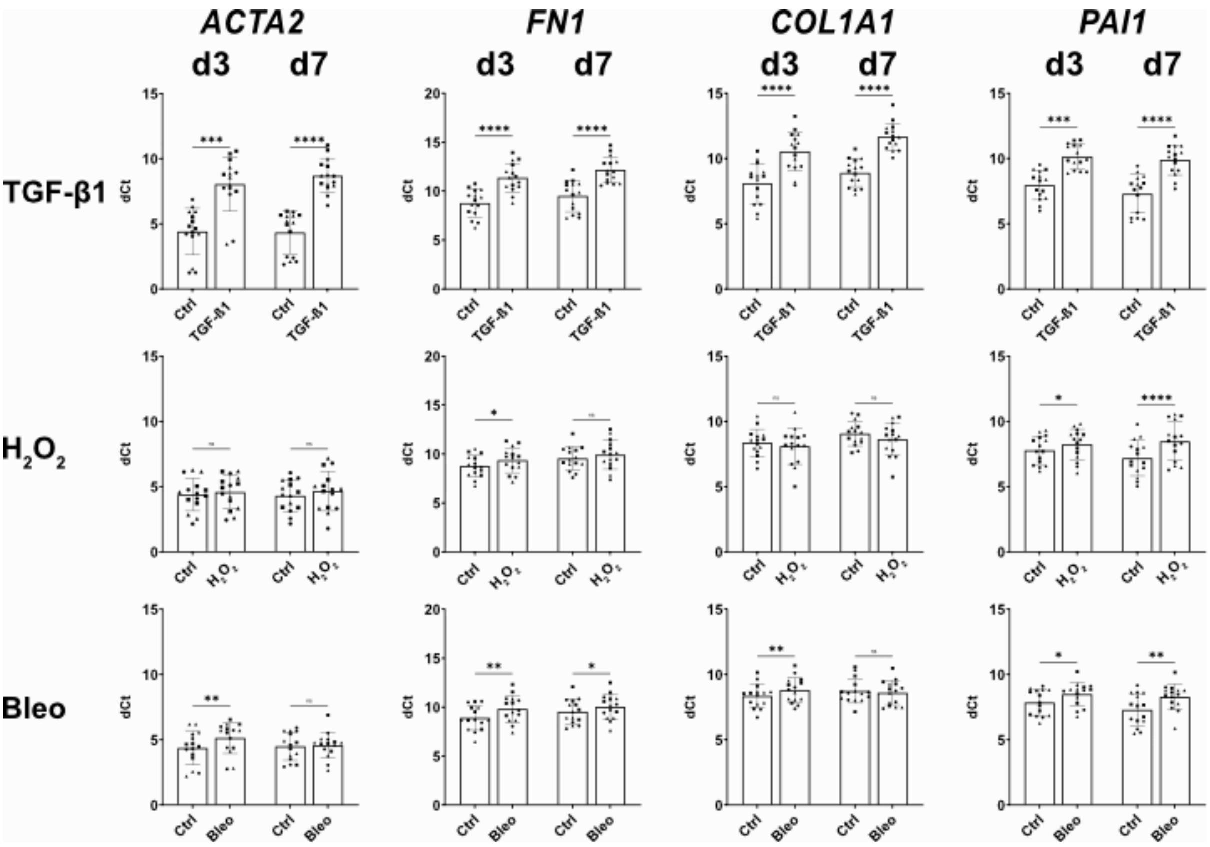


**Supplementary Figure 1. Induction of fibrotic markers in primary human lung fibroblasts.** qRT-PCR to determine the gene expression of fibrosis-related markers after treatment with $H_2O_2$, bleomycin, and TGF-β1 for 3 and 7 days. Data points represent biological replicates from donor (square), IPF (circle), and COPD (triangle). *p-value<0.05 based on paired-Friedman T test.
